# Multiview-SPIM-*µ*PIV for mapping 3C-3D blood flow within the beating zebrafish heart

**DOI:** 10.64898/2026.08.17.745192

**Authors:** J. Jiang, K. Ross, J. M. Taylor

## Abstract

Cardiac blood flow is a regulator of several important developmental and remodelling processes in the heart, including through fluid shear forces sensed by the endothelial cells lining the heart. However, optically mapping these flow fields in the complex 3D geometry of the heart is challenging even in transparent animal models such as the zebrafish. One of the main challenges is the difficulty in measuring the out-of-plane (axial) velocity component, preventing accurate mapping of the complete 3-component-3-dimension (3C-3D) blood flow velocity field; image-based techniques such as microscopic particle image velocimetry (*µ*PIV) traditionally only provide the in-plane flow components. Here we present a computational approach to achieve full time-varying 3C-3D blood flow vector mapping using a standard selective plane illumination microscope (SPIM), based on robust cardiac phase assignment, precise measurement-driven registration of sequentially acquired z-stacks, and PIV data fusion from multiple sample orientations. Our approach holds the key to understanding the complex dynamic flow fields within the developing heart, and their role in shaping cardiac development.

## 1. Introduction

Blood flow exerts shear forces on the surrounding cardiovascular tissue. The key role of cardiac blood flow forces on the development of the embryonic heart in animal models have been extensively studied, especially in zebrafish [1–3]. Accurate mapping of the 3D blood flow vector field is essential for understanding the fluid shear force experienced by the heart endothelium, which provides mechanical cues sensed by receptors such as PIEZO1. Such stimuli are known to be crucial in driving organ development [2, 3] and more local shear-regulated processes such as the formation of the atrial-ventricular valve [4, 5]. Fluid flow mapping also enables quantification of blood flow changes arising from either developmental or disease-driven alterations in cardiac contractility [6] or experimentally induced perturbations [7].

Blood flow in transparent animal models is commonly measured by high speed microscopy of red blood cells. However, blood flow in the heart is a fundamentally 3D process, with the flow vector component perpendicular to the imaging plane making an essential contribution to the flow field and fluid-structure interactions. This third component is difficult to measure experimentally, traditionally requiring stereoscopic methods, volumetric snapshot imaging [8–10] or indirect inference from computational simulations [11]. Although the capability of stereoscopic PIV in measuring all three velocity components of fluid flow has been well established by many studies on a macroscopic scale [12–16], true stereoscopic PIV is rarely used for imaging microscopic fluid flow in live samples. This is because biological imaging requires high-NA objectives to achieve high lateral resolution, but such objectives inherently provide a limited field of view and depth of field. Therefore stereoscopic imaging is in practice difficult to implement in microscopy.

Several alternative *in-vivo* microscopy techniques have been explored with the aim of measuring all three components of blood flow in zebrafish. Truong *et al*. [8], Wang *et al*. [9] and Saberigarakani *et al*. [10] used light field microscopy to visualise blood flow by tracking the motion of individual blood cells. Although they tracked 3D trajectories of individual blood cells, they were not able to scale this approach to map the full time-dependent velocity vector field of blood flow throughout the whole heart. Holographic microscopy [17] is another viable method for measuring 3-component blood flow by reconstructing the instantaneous positions of blood cells in 3D. However, both light field and holographic microscopy trade resolution, especially in the axial direction, for their capability of acquiring volumetric snapshots, as well as suffering from reconstruction artefacts [8] and degraded imaging performance when faced with densely packed blood cells in the heart.

In an alternative approach, Chan *et al*. [18] developed a multi-view strategy to reconstruct the 3D flow field from brightfield images acquired at rotated sample orientations. However, brightfield *µ*PIV suffers from so-called depth of correlation artifacts [19], which are liable to bias the measured velocity values [20] and prevent faithful recovery of the 3D flow field. A solution to the depth of correlation issue is to use selective plane illumination microscopy (SPIM) for *in-vivo* measurement of fluorescently-labelled blood flow [20], delivering good optical sectioning and low phototoxicity. Zickus *et al*. [20] combined SPIM with micro particle image velocimetry (*µ*PIV) to solve the depth-of-correlation problem – but at the expense of only being able to map the two in-plane velocity components of cardiac blood flow in 3 spatial dimensions (2C-3D). Zhang *et al*. [21] subsequently adopted a similar approach for imaging blood flow in micro-vessels. They used a dual-mode light sheet microscope to infer the full 3-component velocity of individual blood cells, based on measurements of the 2-component velocity as projected onto the imaging plane. Since the blood flow vectors were known to be confined by the vessel geometry to a (skewed) two-dimensional plane, the 3-component velocity could be inferred after measuring the projection angle from volumetric images of the thin blood vessels. However, this approach cannot be extended to measuring the full 3-component 3-dimensional (3C-3D) blood flow in the heart, since the flow is not confined to a cylindrical lumen. In the heart the *direction* of the blood flow cannot be deduced from the geometry alone, and needs to be measured experimentally.

To overcome these challenges we present a new computational technique capable of mapping the full 3C-3D blood flow vector field within a beating zebrafish heart. Our method relies on the fusion of temporally and spatially registered SPIM images acquired at multiple sample orientations. Our technique follows a similar tomographic principle to the work of Chan *et al*. [18], but our method takes a different approach to 3D velocity calculation, introduces a new strategy for measurement-driven registration and uncertainty quantification, and benefits from the good optical sectioning capability of the SPIM to avoid depth of correlation artifacts in the measurements.

## 2. Method

### 2.1. Sample preparation and imaging

We imaged a Tg(gata1:dsRed; fli1:GFP) zebrafish line, giving red fluorescent labelling of red blood cells and green fluorescent labelling of cardiac endothelial cells. Zebrafish were incubated with 0.002% phenylthiourea (Sigma-Aldrich, P7629-25G) to reduce the formation of dark pigments on the skin for better optical access [22]. A 4.5 days post-fertilization zebrafish was sedated with tricaine (Thermo Scientific, 118000100) with a concentration of 600 µM (170µg mL^−1^), and embedded live in 1% low melting-point agarose (Sigma-Aldrich, A9414-25G) to ensure sample stability. The zebrafish was mounted in fluorinated ethylene propylene (FEP) [23] tubing (Adtech, STW17) for refractive index matching to water, attached to an 18 gauge blunt cannula (e.g., Farnell 918050-TE) on a syringe mount (Becton Dickinson, BD303172) and inserted into the rotational stage of the SPIM system shown in Fig. 1.

**Fig. 1.**
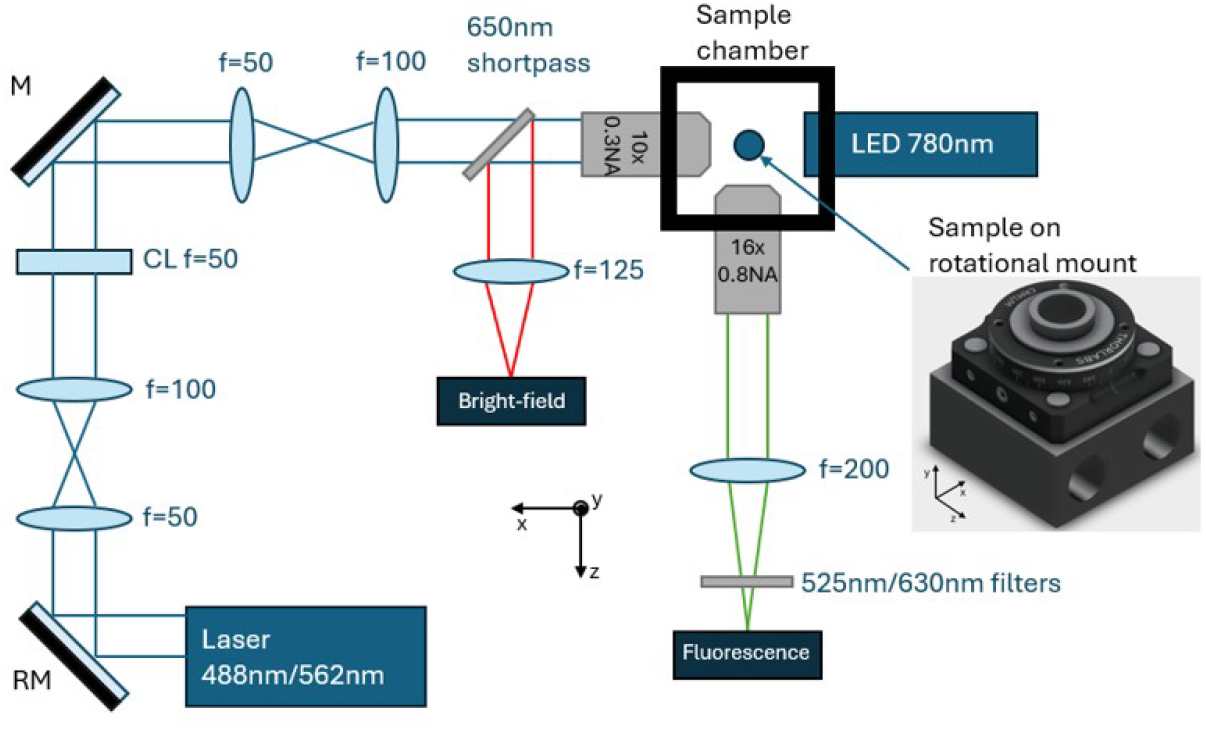
SPIM setup for the simultaneous imaging of zebrafish in brightfield and fluorescent channels. The 525/39nm emission filter is used with 488nm laser excitation to image GFP fluorophore in heart endothelium, and the filter can be swapped to 630/69nm to image dsRed fluorophore in red blood cells with 562nm laser excitation. A resonant mirror (RM) is used to mitigate shadow effects in images caused by dark pigments in zebrafish skin. The sample is mounted in a syringe, inserted into the rotational mount shown. The sample can be rotated around the y axis using the rotation plate. The mount is secured on a motorised stage that can move along three axes.

### 2.2. Multiview z-stack acquisition

Images of the GFP fluorophore were acquired to visualise the heart structure and to perform multi-view stack registration. In order to measure the blood flow velocity, we use the dsRed-labelled red blood cells as tracer particles. Images of red blood cells were acquired and the flow velocity was measured by particle image velocimetry (Section 2.3).

Since the structure of the heart and the blood flow change across the periodic heartbeat, a retrospective optical gating technique [24] was used to assess the heart phase and assign each acquired fluorescent image an associated phase value between 0−2*π*. Images acquired at different axial locations were grouped into phase bins. This effectively achieves volumetric imaging across the heartbeat, allowing a consistent 3D representation of the heart’s structure to be reconstructed at any phase in the heart cycle, and allowing sets of blood flow image pairs to be identified where each frame pair corresponds to the same phase in the heart cycle.

Our optical gating technique exploits the periodic change in the heart shape as observed in a brightfield channel captured simultaneous to all fluorescence imaging [25]. For every brightfield image, the phase of the heart was determined against a reference sequence of the heartbeat – and then comparison and interpolation between frame timestamps allowed the phases of fluorescence frames to be determined from the known phases of nearby brightfield frames. All the measured phase values are defined relative to the initially-identified reference heartbeat sequence. Since rotating the sample changes the appearance of the heart as viewed in the brightfield channel, a new reference sequence is required for each sample orientation, and phase values measured for each individual sample orientation have an *a priori* unknown fixed phase shift between them. This phase shift must be computationally determined as part of the registration process (Sections 2.6 & 2.7).

The image acquisition procedure is represented in Fig. 2. Fluorescence images were acquired using a CMOS camera (Ximea MC031MG-SY-UB). For each sample orientation, sequential z-stacks for the green and red channels were acquired by translating the sample stage along the z-axis. Each z-stack contains 31 planes with a 4µm plane separation, ensuring the image acquisition time is minimised while retaining sufficient axial resolution of the resulting flow field measurement. To minimise phototoxicity the excitation laser was only triggered for 1 ms per frame. In standard pulsed mode, frames are captured at regular intervals with the laser on throughout each frame; in PIV mode, the laser is triggered near the end of one frame exposure, and at the start of the subsequent frame exposure, with the temporal separation between laser pulses being 1 ms. This technique increases the temporal resolution of image pairs beyond the maximum achievable camera frame rate [20]. Each plane in the red channel was imaged in PIV mode over the course of 18 seconds at 200 frames per second, to obtain at least 50 frames per 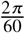 phase bin. This ensures a sufficient number of frames for correlation-averaged PIV. For the green channel, each plane was imaged in pulsed mode over the course of 2 seconds at 200 frames per second to ensure sufficiently dense sampling of images across the 0 − 2*π* phase range. To achieve an isotropic voxel aspect ratio for subsequent processing, the green channel z-stacks were upsampled along the z-axis using linear spline interpolation (Scipy [26]). This upsampling helps with multi-view stack registration and visualisation of the orthogonal cross sections of the heart structure. The complete image dataset is available in the data repository [27] and the Python scripts used for our analysis are available on GitHub [28].

**Fig. 2.**
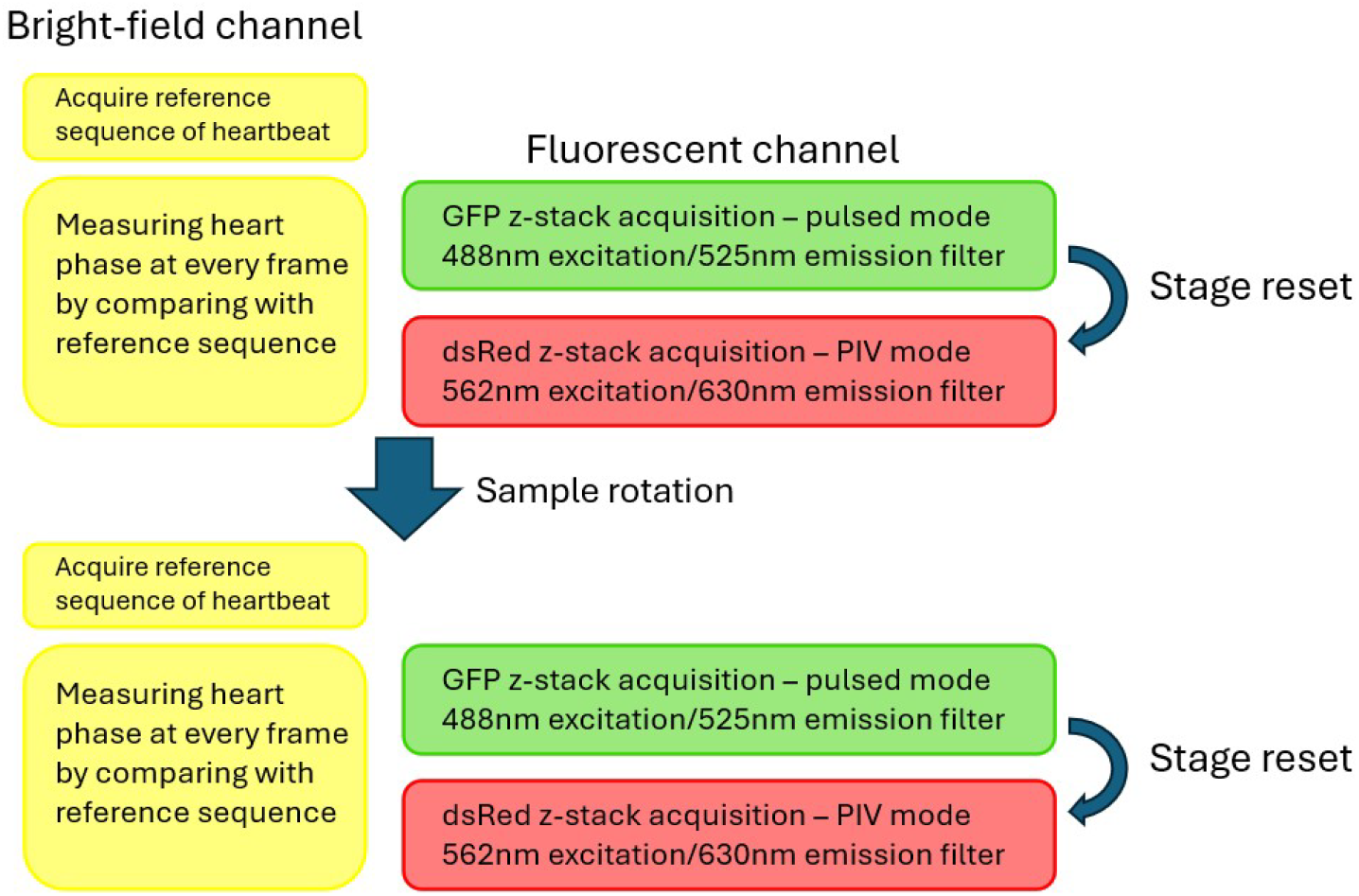
Schematic illustration of the imaging procedures. Z-stacks were acquired by translating the rotational mount, shown in Fig. 1, along the z-axis. The sample was imaged simultaneously in the brightfield channel for providing phase information of the heart.

### 2.3. Correlation-averaged PIV

The flow velocity was calculated by measuring the displacement of red blood cells between two images, in a process known as cross-correlation PIV [29]. The process of cross-correlation is illustrated in Fig. 3. A correlation matrix is calculated between an interrogation window (IW) taken from the first image and a search range from the second image. We calculated the correlation product by taking the sum of the pixel-wise absolute differences (SAD) between two windows. The average of multiple correlation matrices, calculated from multiple image pairs within the same phase bin, was taken for *correlation averaging* (see below). The overall process can be mathematically expressed as:

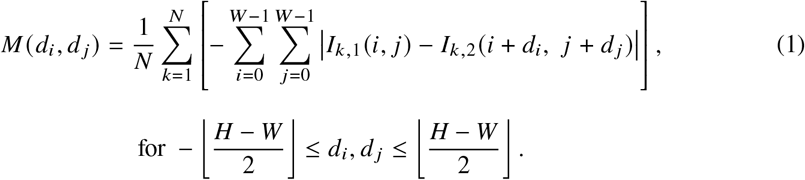

where *M d*_*i*_, *d* _*j*_ is an element of the correlation matrix, averaged from *N* independent correlation matrices; *W* is the size of the IW, *H* is the size of the search range, *I*_*k*,1_ and *I*_*k*,2_ are pixel values of two images of pair *k*, and *d*_*i*_, *d* _*j*_ is the IW shift in the x and y directions. Conventionally a lower SAD value corresponds to a lower difference (higher similarity) between windows. However, note that we use negative SAD values such that the peak of the correlation matrix corresponds to the highest similarity, for direct compatibility with Gaussian peak-finding algorithms (OpenPIV [30]). The optimal IW shift is found by locating the peak of the correlation matrix to sub-pixel precision. The flow velocity at the position of the IW is then calculated from the measured IW shift and the known temporal separation between images:

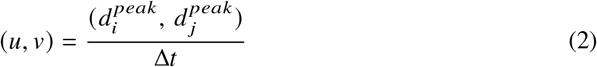

where (*u, v*) is the calculated velocity in x and y, 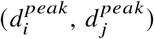 is the feature displacement in x and y measured as the peak position of the correlation matrix and Δ*t* is the temporal separation between frames. For this experiment Δ*t*=2ms, the temporal separation between laser pulses.

**Fig. 3.**
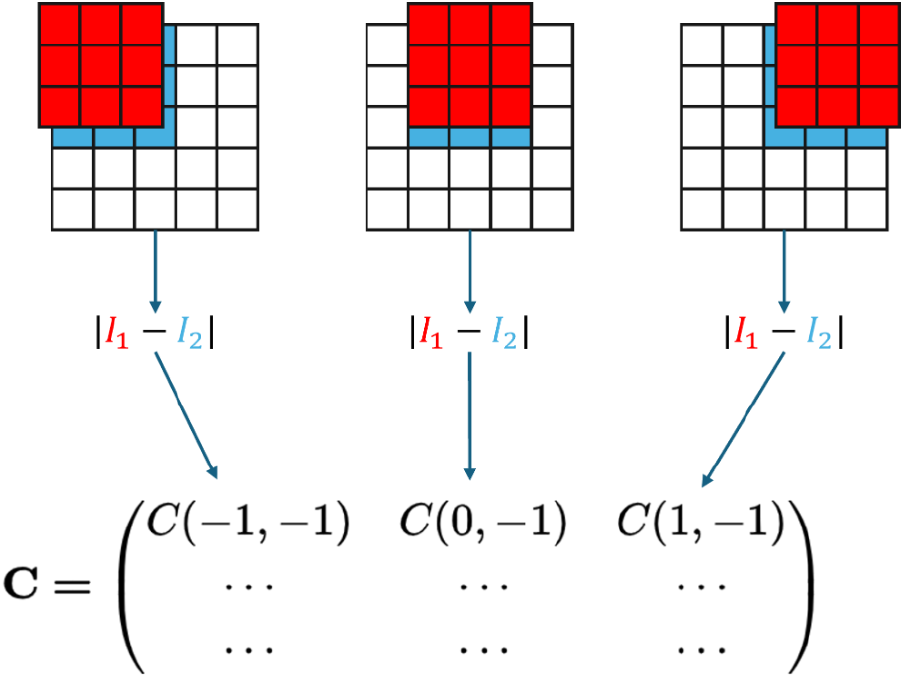
Image cross-correlation for flow measurement. In this reduced-size example, a 3×3 interrogation window (IW) shown in red shifts around a 5×5 search range. The sum of absolute differences (SAD) between the pixels of interrogation window and corresponding pixels in the search range (blue) is calculated, resulting in a correlation matrix, **C**, where the displacement of features in the IW is given by the peak position of the correlation matrix.

Correlation-averaging enhances the signal-to-noise ratio of the correlation matrix by amplifying the peak signal. This is required when the seeding density and the image signal-to-noise ratio is low [31]. This is common when imaging in the heart, as the blood is not saturated with blood cells and a single IW may contain few or no clear features from blood cells. To enhance the signal-to-noise ratio, separate correlation matrices were calculated between multiple frame pairs in the same phase bin, based on the reasonable assumption that the blood flow is consistent at the same heart phase across different heartbeats. The feature displacement was then calculated from the averaged correlation matrix.

### 2.4. Multiview velocity fusion

As Zhang *et al*. [21] demonstrated in their studies, the velocity of the blood flow measured from 2D images is the projection of the true 3D flow velocity onto the imaging plane [20]. Therefore a different projection of the 3D flow vector can be obtained by rotating the sample. At any given position in the heart, the blood flow vector can be expressed as: ***L*** = (*u, v, w*); rotating the sample around the y-axis by an angle *γ* results in a rotated vector: ***L*′** = (*u*′, *v*′, *w*′). Applying the standard rotation matrix ***R***_*y*_ (*γ*), the rotated vector can be expressed as:

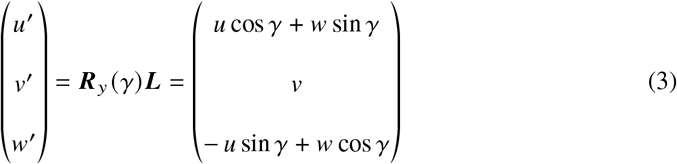

which can be rearranged to give the z-component of the flow vector:

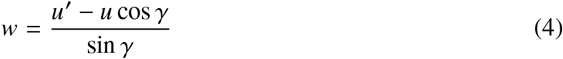

where *u* and *u*′ represent the x component of the velocity as measured by PIV in the two different sample orientations.

It is important that both *u* and *u*′ are measured at the exact same positions in the heart, so they capture the projections of the same blood flow vector. To measure *u* and *u*′, PIV analysis was performed at the precise intersection positions between the two z-stacks as shown in Fig. 4. Since the sample could not be mounted precisely on the rotation axis of the stage, the sample required repositioning after the rotation. Therefore image stacks acquired before and after repositioning the sample need to be registered. We developed a rigorous and robust registration process to reach pixel level accuracy in stack registration. The relative intersection positions were calculated after stack registration based on the green channel, and PIV was performed on red channel images to obtain *u* and *u*′, thereby providing all three flow components at those positions in the heart.

**Fig. 4.**
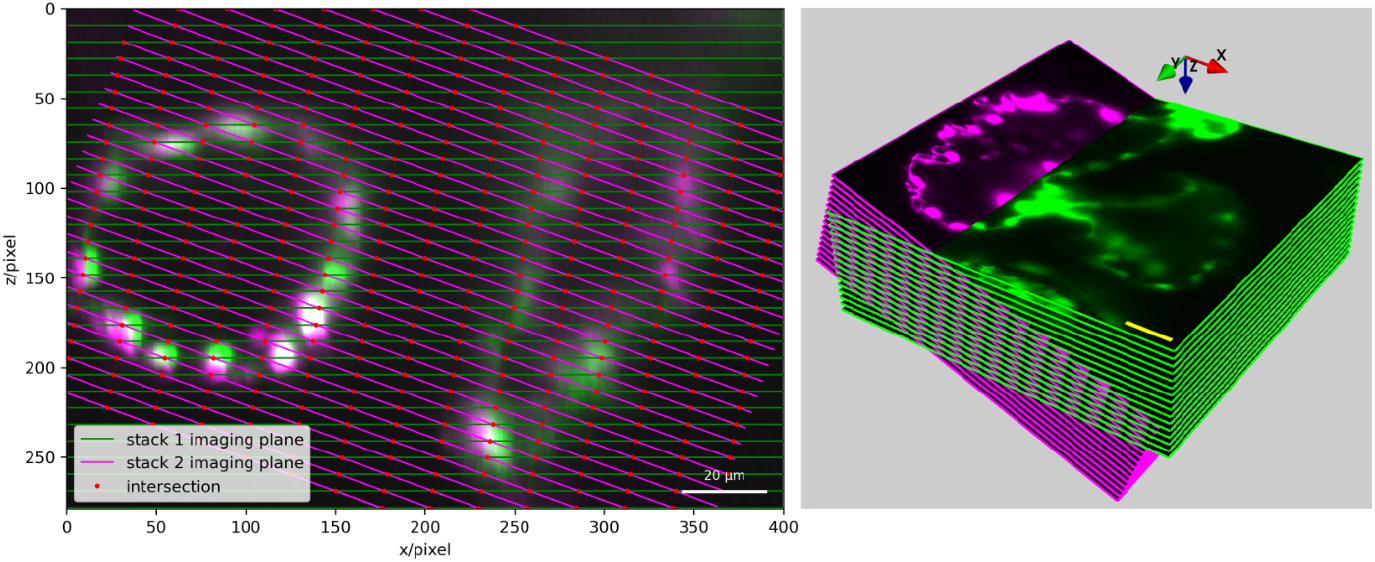
Left: 2D illustration of intersection between two stacks acquired at first sample orientation (green) and at rotated sample orientation (magenta), images shown are XZ cross sections of the heart z-stacks. Red points label the intersection positions where the blood flow was measured. Right: 3D illustration of intersection between two stacks. Stacks are partially shown for better visibility of the central cross section of the heart. Scale bar 20 µm.

### 2.5. Estimation of PIV uncertainty

As mentioned in Section 2.3, correlation averaging is a method to fuse information obtained from multiple frame-pairs acquired from different heartbeats. Although it is reasonable to assume consistent blood flow between different heartbeats, relying on the periodicity of the heart, it is important to verify this consistency and quantify the small variations in blood flow within the same phase bin. This provides quantitative estimates of uncertainties in our measurements, allowing us to evaluate our approach. Additionally, knowing the uncertainties in correlation averaged PIV enables us to optimise the *stack registration* (see Section 2.7).

No well-established, robust method exists to determine the uncertainty in the result of PIV correlation averaging, as the calculation yields only a single value and the variance cannot be explicitly calculated. Velocity vectors from single isolated frame pairs of flowing blood are not a reliable basis for statistical analysis since, as discussed earlier, some frame pairs contain little or no information on the flow. Therefore we developed a new (to the best of our knowledge) approach to estimate PIV uncertainty *via* bootstrapping. Bootstrapping is a statistical method first proposed by Efron [32], where the mean and variance of a distribution can be estimated by repeated resampling of the distribution. In our case, to estimate PIV uncertainty, frame pairs were resampled with replacement. Correlation averaged PIV was performed on each resampled subset of frame pairs. Measurements from all subsets then form a distribution from which the variance was measured. Fig. 5 shows an example of bootstrapped PIV measurement. By resampling the subset of frame pairs used in correlation averaging for a large number of iterations (50000 in this example), the result of the PIV measurements approximates a normal distribution, where the standard deviation gives the uncertainty of PIV measurement.

**Fig. 5.**
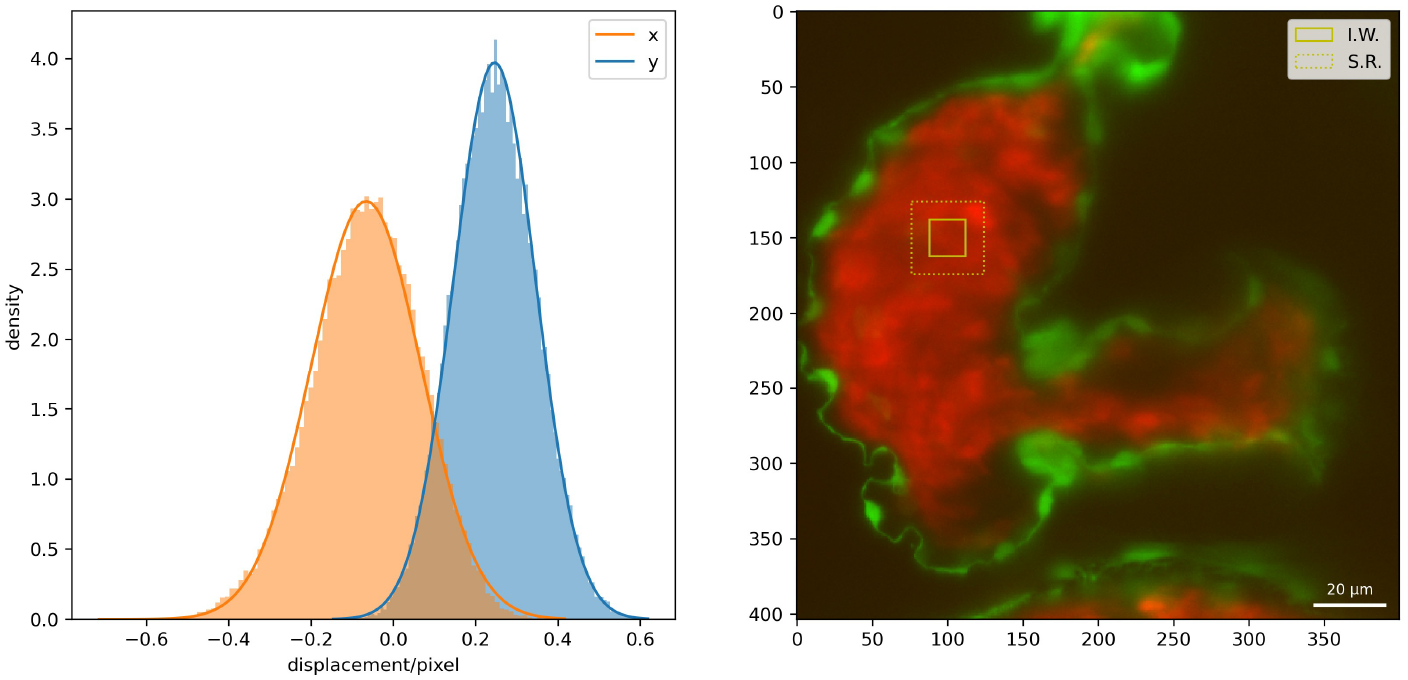
Left: distribution of the feature displacement in x and y measured by bootstrapped correlation averaged PIV. Displacement in x follows a normal distribution with mean −0.066 and standard deviation 0.13; displacement in y also follows a normal distribution with mean 0.25 and standard deviation 0.10. Right: superimposed image of the cardiac endothelium and red blood cells. Yellow squares mark the interrogation window and the search range used in PIV.

Recalling Eqn. 4, the uncertainty in the calculated axial velocity, *w*, is given by:

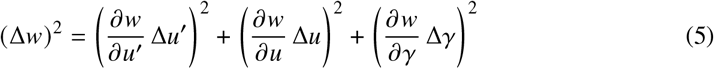

where *u*′ and *u* are the x component of the velocity as measured by PIV in the two different sample orientations and *γ* is the rotation angle between sample orientations. Solving the partial derivatives gives:

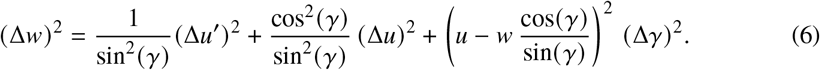

### 2.6. SSIM-based stack registration

As explained in Section 2.4, z-stacks acquired at each sample orientation need to be registered to yield the correct intersection positions. Registration involves a spatial transformation (rotation by Δ*θ*, shift by Δ*x*, Δ*y*, Δ*z*) and a phase shift (Δ*ϕ*). We performed an initial registration estimate based on the structural similarity index measurement (SSIM) between the green channel images of the heart endothelium acquired at each sample orientation. As illustrated in Fig. 6, we applied a phase shift and an affine transform to the first z-stack to account for the rotation and translation of the heart during the rotation and repositioning of the sample. We then used the SSIM between two stacks to optimise all five registration parameters. To minimise the computation time, phase registration was performed independently using a set of visually estimated spatial registration parameters. Δ*ϕ* was then optimised by varying it between 0-2*π* to find the SSIM maximum. After applying the optimised Δ*ϕ* to the z-stack, Δ*θ*, Δ*x*, Δ*y* and Δ*z* were simultaneously optimised using a Bayesian optimiser from Scikit [33], chosen for its efficiency at optimising expensive multivariate functions. We note that the optimisation yields slightly different results at different phases of the heartbeat. Therefore we performed optimisation at intervals of 2*π*/30 across the full heartbeat, and obtained the average and standard deviation of each stack registration parameter. These parameters were used to calculate the intersection positions, as illustrated in Fig. 4, where the flow velocity was measured using PIV.

**Fig. 6.**
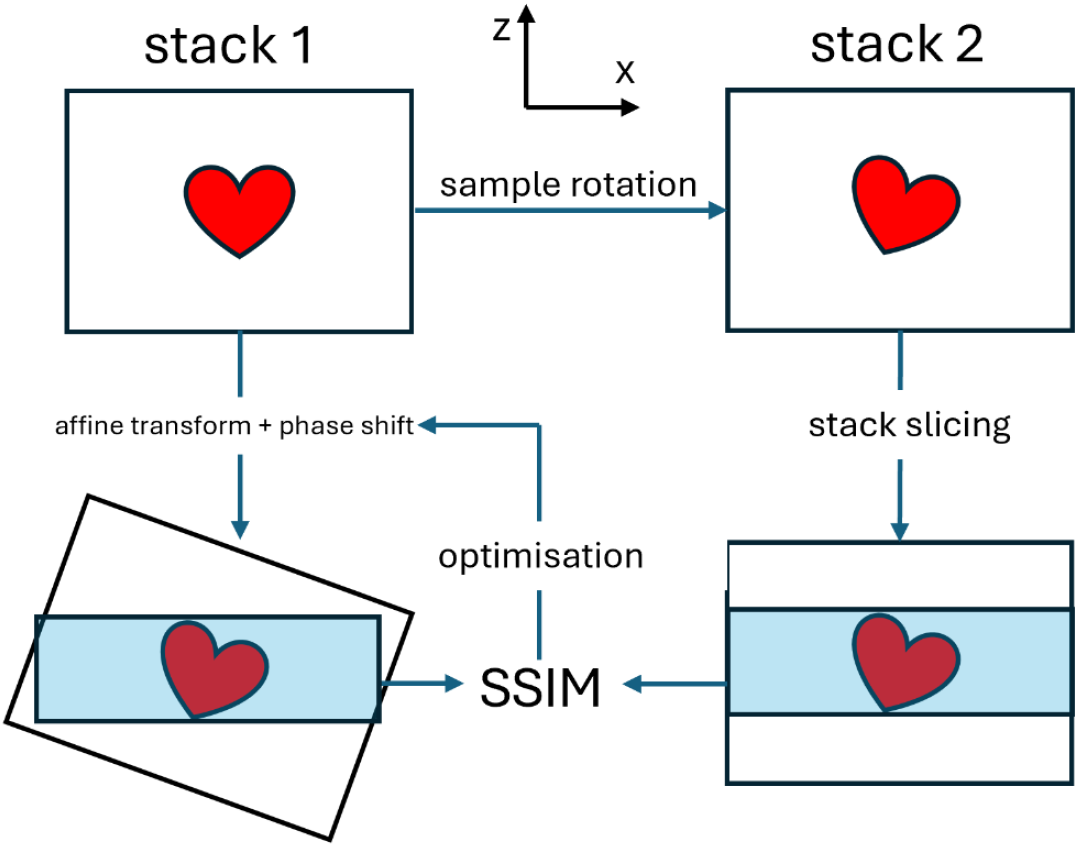
Schematic of the stack registration process. The red heart represents the orientation of the zebrafish heart viewed in the x-z plane. The heart was physically rotated around the y axis between the acquisition of two z-stacks. An affine transform (rotation by Δ*θ* and shift by Δ*x*, Δ*y*, Δ*z*) and a phase shift of Δ*ϕ* was applied to the first upsampled green channel z-stack, and a structural similarity index measurement (SSIM) was calculated between the central cyan-shaded part of two stacks. Stacks were sliced to avoid pixels padded during the affine transform as the padding value would bias the SSIM calculation. The SSIM value was used to optimise all five registration parameters, which were then used to accurately calculate the exact intersection positions between two stacks.

### 2.7. Measurement-driven stack registration and validation

An important principle of our multi-view velocity fusion is that we are fusing two projections of the *same* blood flow vector, acquired from two sample orientations. Accurate stack registration is therefore essential. To optimise and quantify this stack registration in the red imaging channel (blood cell velocity), we compared the y-velocities at the intersection positions as determined from the SSIM-based stack registration. The sample was rotated around the y-axis; therefore the velocity component parallel to the y-axis should be rotationally invariant. Fig. 7 compares the 2D velocity fields and the y-velocities calculated from each of the two stacks, at the same co-registered positions in the heart. Perfect rotational invariance would result in a diagonal line with equal y-velocities at any position in the heart, as represented by the red line in Fig. 7 right. The agreement of y-velocities between two stacks was quantified using the reduced *χ*^2^ value. A main source of deviations from the diagonal line is errors in stack registration; we observe that perturbing the stack registration has a prominent impact on the *χ*^2^ value, with some perturbations giving an improved *χ*^2^ value compared to that obtained following SSIM registration. This observation allowed us to perform stack registration based on the flow velocity measurement, by optimising stack registration parameters to yield the minimum *χ*^2^ value.

**Fig. 7.**
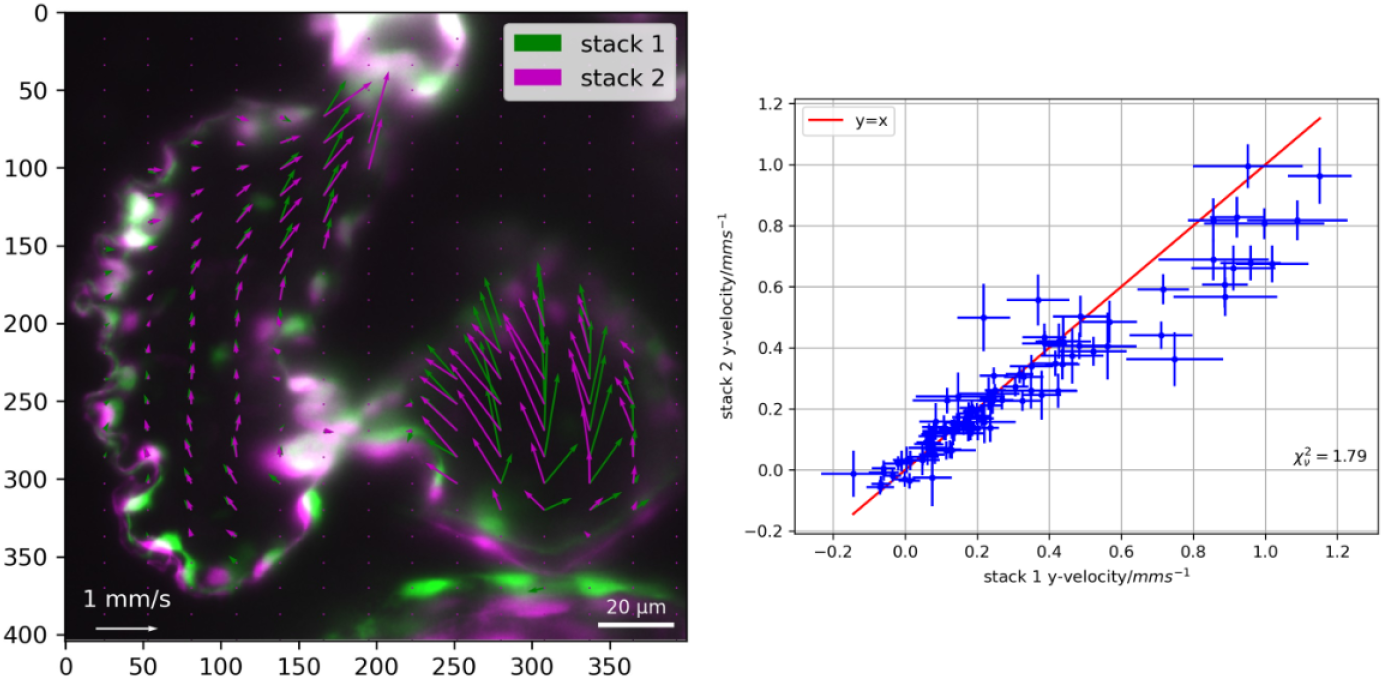
Left: cardiac blood flow vector field superimposed on the GFP images from two z-stacks at two sample orientations. Flow velocity measured by PIV, using dsRed images from two z-stacks; each vector represents one co-registered location in the heart. The horizontal (x) component of these vectors give *u* and *u*′ values that are used to calculate the flow velocity perpendicular to the imaging plane. The GFP image shown in green is the central plane of stack 1, after affine transform is applied, whereas the GFP image shown in magenta is the central plane of stack 2. Right: y-velocities of the vector fields shown on the left. Uncertainties given by the standard deviation of the bootstrapped PIV distribution. A rotational-invariant vector field would result in the same y-velocities, indicated by the red diagonal line. Agreement between the measured y-velocities was statistically quantified using the reduced *χ*^2^ value which is 1.79 for this plot.

Fig. 8 shows the variation in the reduced *χ*^2^ value as a function of the stack registration parameters. The reduced *χ*^2^ value increases as parameters move away from the minimum, representing increasing deviations from rotational invariance. Optimisation was performed using an iterative Gauss-Seidel method, where parameters were optimised sequentially with the initial values set as the SSIM optimum. Ventricular flow measurements in three z-planes (10th, 15th and 20th planes) were used to sample measurements throughout the whole heart, while maintaining a reasonable computation time. The minima of the *χ*^2^ curves (green lines in Fig. 8) were calculated using local parabolic interpolation. Although this measurement-driven approach reached sub-pixel precision in stack registration, it is time-consuming due to the large number of PIV measurements required. We therefore rely on the SSIM-optimisation to provide good initial values, such that only two iterations are required for the Gauss-Seidel method to converge. The combination of image-based SSIM-optimisation and measurement-driven *χ*^2^ optimisation ensures efficient and precise registration of multi-view image stacks.

**Fig. 8.**
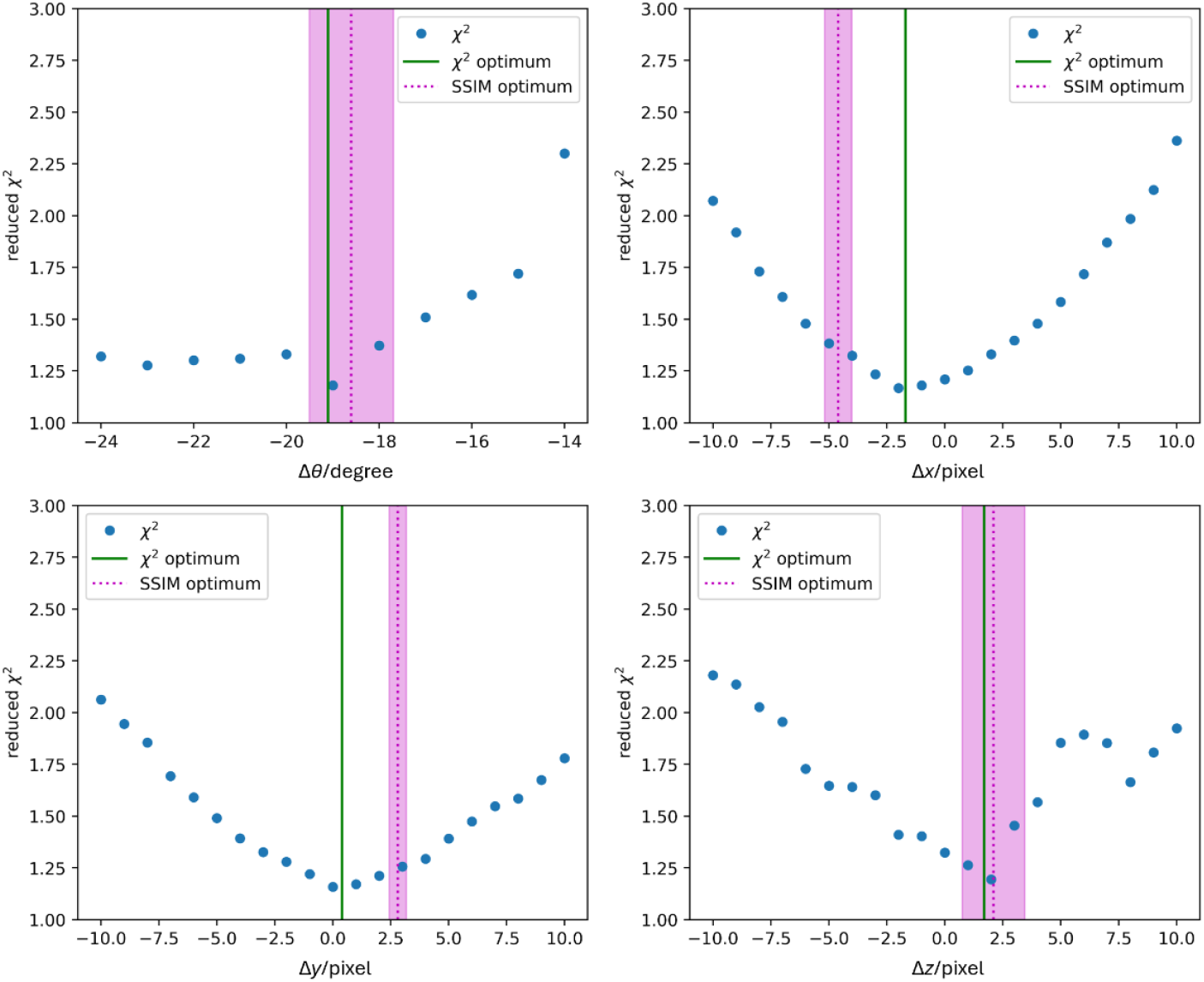
Reduced *χ*^2^ values of y-velocities measured from two stacks, varied with stack registration parameters. Δ*θ* has a 1 degree interval, while Δ*x*, Δ*y*, Δ*z* have a 1 pixel interval. A lower *χ*^2^ value indicates better stack registration, because the agreement between y-velocities supports the rotational-invariant assumption. Each plot depicts one parameter being varied with the other three held constant. The dotted magenta lines show the optimal stack registration parameters obtained from the SSIM Bayesian optimisation, and the shaded regions represent the standard deviations in the parameters obtained at different phase of the heartbeat. Green lines show the optimal parameters for flow-based registration.

## 3. Flow field mapping and discussion

### 3.1. 3C-3D blood flow vector field

Fig. 9 shows the 3C-3D blood flow vector field as determined at two different phases of the heartbeat. GFP cross sections of the heart endothelium (wall lining) are overlaid to show the relative position of the blood flow within the heart chambers. We observe in Fig. 9a and Fig. 9b that there is a strong flow entering the atrium from the sinus venosus (major vein leading into the heart), through the sinoatrial node. The orientation of the vessel is approximately orthogonal to the imaging plane; thus the high flow velocity in the z direction has been successfully recovered using our multi-view fusion technique. We also observe in Fig. 9d and Fig. 9f successful recovery of the expected convergent flow field in the z direction as the blood approaches the atrial-ventricular canal (AVC; the junction between the atrium and the ventricle). The z-velocity of the flow increases as it approaches the AVC due to the narrowing constriction. Fig. 10 and Fig. 11 show the projections of the 3D vector field on the three cross sections shown in Fig. 9, for a clearer view of the vector field. Note the axial convergence of the blood flow, most clearly visible in the XZ plane of Fig. 11; and the axial perturbation to the flow caused by the bulging heart tissue visible in the YZ plane. These examples illustrate how the axial velocities obtained from our multiview fusion accurately reflect the expected flow behaviour dictated by fundamental fluid-dynamic principles.

**Fig. 9.**
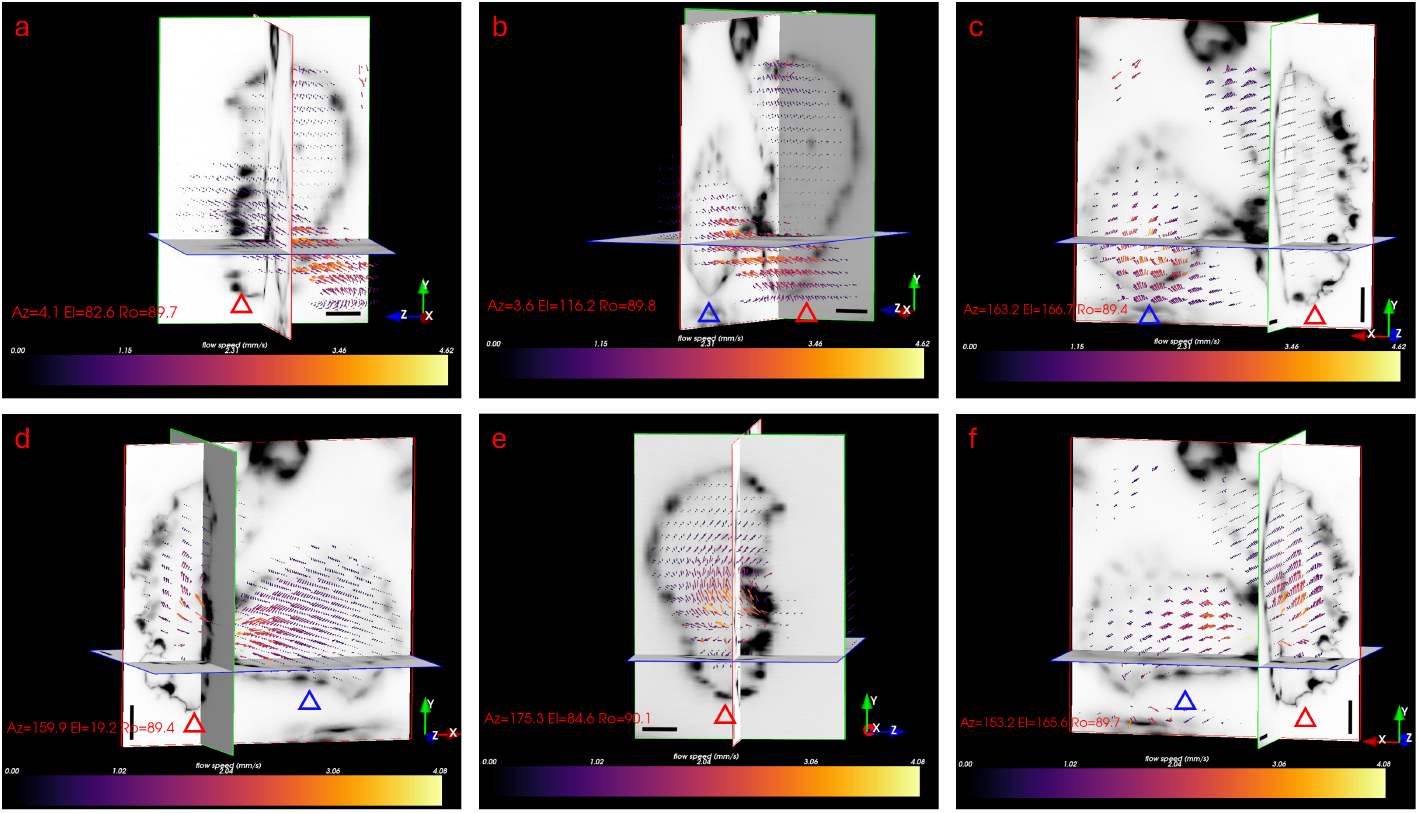
a,b,c: orthographic views of the 3D vector field snapshot at the diastolic phase of the heartbeat; d,e,f: snapshot at the systolic phase of the heartbeat. Vector field superimposed on the XY cross section (red border), YZ cross section (green border) and XZ cross section (blue border) of the upsampled green channel z-stack. Flow velocity in the z-axis was calculated from multiview velocity fusion. To facilitate orientation within images, blue triangles label the atrium, and red triangles label the ventricle. Scale bar 20 *µ*m. Images rendered in Mayavi [34]. See Visualisation 1 and 2 for a complete view of the vector field.

**Fig. 10.**
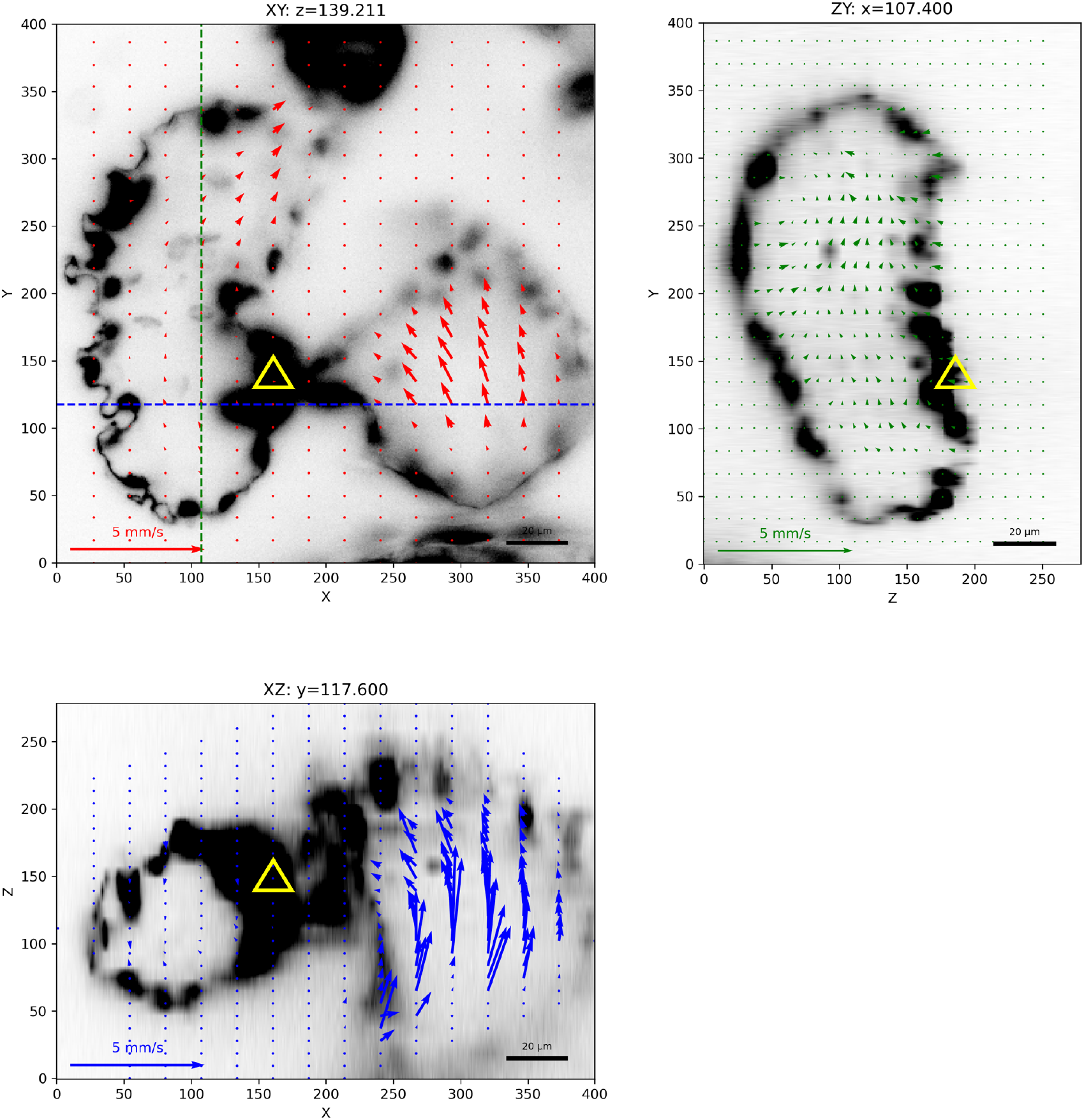
Projections of the 3C-3D vector field on the three orthogonal cross sections shown in Fig. 9(a,b,c). The flow was measured during the diastolic phase of the heartbeat, when a strong influx of blood from the sinus venosus is induced by the relaxation of the atrium. Images upscaled in the z-axis to maintain a consistent aspect ratio. Yellow triangles mark the approximate position of the AVC. Black-level of images adjusted to enhance image contrast for improved visibility of the heart wall.

**Fig. 11.**
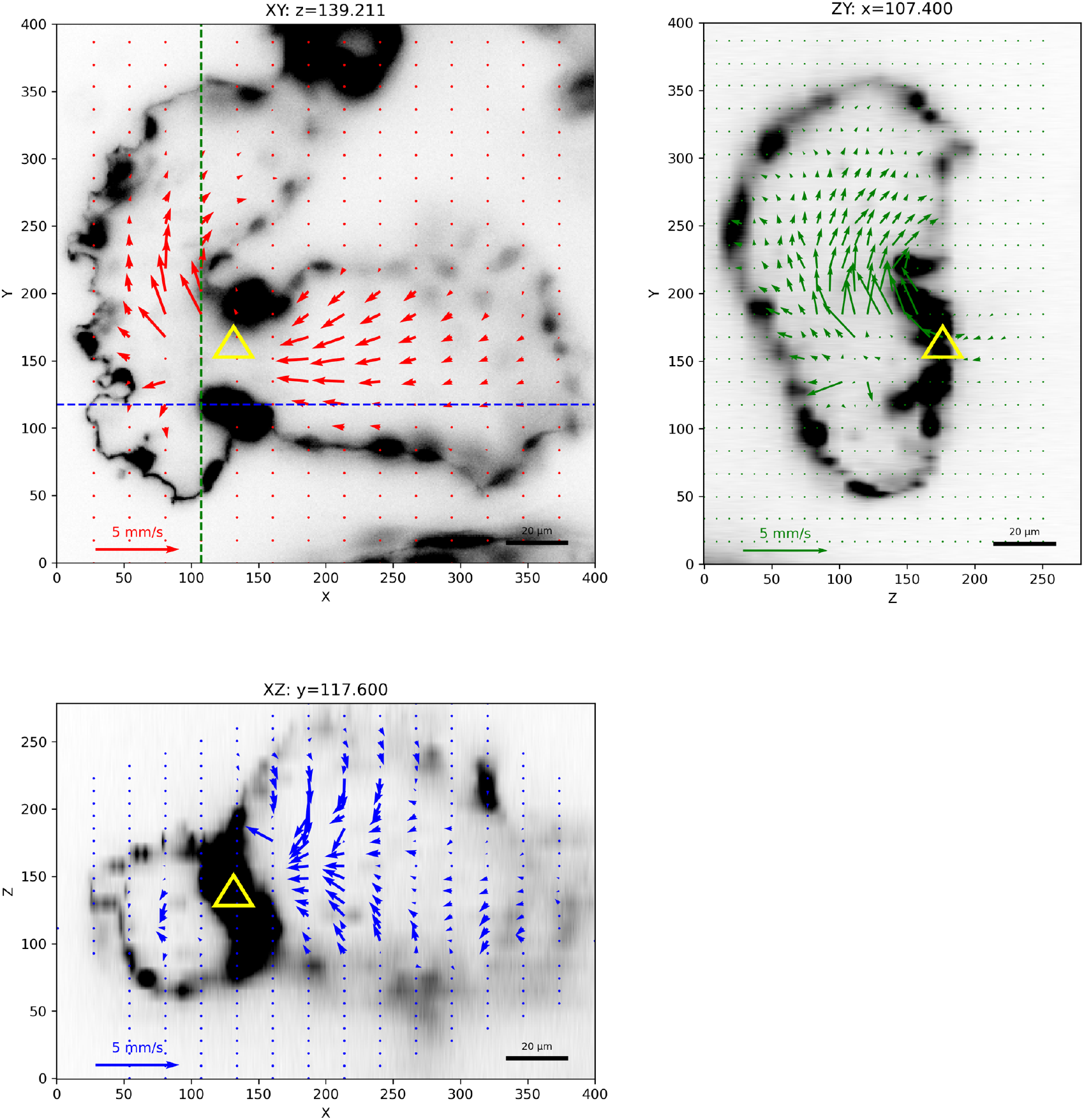
Projections of the 3C-3D vector field on the three orthogonal cross sections shown in Fig. 9(d,e,f). The flow was measured during the start of the systolic phase of the heartbeat. We observe a strong directional flow from the atrium towards the ventricle, across the AVC marked by yellow triangles. Images upscaled in the z-axis to maintain a consistent aspect ratio. Black-level of images adjusted to enhance image contrast for improved visibility of the heart wall. Note the region within the AVC did not yield any valid measurements, as PIV performs poorly in regions with extreme flow velocities that induce transformations of the imaged features beyond linear translation (e.g. rotation and deformation of blood cells).

These observations show that there is a significant component of the blood flow in the direction perpendicular to the imaging plane, particularly near the entrance (sinus venosus) and the exit (AVC) of the atrium. The axial flow contributes significantly to the shear force between the blood flow and the heart tissue. Due to the looped architecture of the heart, some regions of the flow field will *always* be axially-dominated, regardless of the sample orientation. It is therefore crucial to measure the full 3C-3D flow to determine the true shear forces within the heart.

### 3.2. PIV resolution

Since PIV measures feature displacement by cross-correlating an IW against a wider search area, the result will be (approximately speaking) an average of the feature displacement field across the IW. The size of the IW needs to be sufficiently large to capture enough tracers for ample signal-to-noise ratio, while an overly-large IW entails excessively spatial averaging and hence reduces effective spatial resolution. The resolution of PIV measurement is on the same order of magnitude as the size of IW, which is 24×24 pixels (1 pixel = 0.431*µm*). We note that despite this large IW size and limited resolution, PIV measurements are still sensitive to pixel-wise changes in the IW position (analogous to the distinction between localization and resolution when imaging individual particles). Therefore, as demonstrated in Section 2.7, it was essential that the positions of individual PIV measurements were correctly matched between different 3D sample views with pixel-level accuracy.

### 3.3. Inter-beat blood flow variability

Correlation averaging improves flow vector signal-to-noise ratio by combining images of red blood cells acquired at the same phase across different heartbeats [20]. We used a sophisticated cardiac phase measurement approach described in Section 2.2 to ensure all images used in correlation averaging lie within the same 2*π*/60 phase bin. Nevertheless, the pulsatile, constantly-accelerating flow within the heart means that there will still be a small quasi-linear variation in local blood flow velocity across a phase bin, in addition to small variability in heart rhythm from one beat to another [35]. We quantified the effect of these variations, along with other sources of experimental error, using a novel bootstrapping strategy (Section 2.5). In principle higher accuracy could be achieved in our velocity measurements by narrowing the phase bins. However, this would require a longer imaging time to acquire sufficient images in each phase bin, so for any given experiment a balance must be struck between accuracy and a longer-duration experiment increasing the susceptibility to long-term variations in the blood flow due e.g. to development, and phototoxic damage to the heart as a consequence of increased imaging duration.

Over the course of the imaging experiment we did observe a reduction in heart rate from an average of 1.82 Hz when imaging the first sample orientation to an average of 1.73 Hz for the second orientation. This may have caused blood flow to be generally slower during the acquisition of the second z-stack; signs of this are visible in Fig. 7 right, where the y-velocity measured from stack 1 can be seen to be slightly higher in the regions of greatest flow velocity. This decline in heart rate is likely caused by a combination of imaging at room temperature (rather than the 28°C of the incubator), chemical sedation, and potentially accumulated laser exposure. We attempted to keep the heart rate more constant by adjusting the sedative dosage, allowing time for the zebrafish to stabilise before imaging, and minimising the imaging time. More work is required to understand more closely the exact causes of the changing heart rate and maintain completely constant blood flow conditions throughout imaging.

### 3.4. Two-phase blood flow

Most non-invasive experiments use blood cells as tracer particles to measure the velocity of blood flow, since the flow of the (featureless) blood plasma itself cannot easily be directly tracked. The ability to measure the flow profile near the wall of the heart/blood vessel is of particular interest for the measurement of shear force. We measured the blood flow in the heart by tracking the motion of blood cells, but high-precision measurements of shear force are a particularly challenging application. Near the edges of a chamber the blood cell density is lower as a consequence of the Fåhræus–Lindqvist effect [36], as can be observed in Fig. 5 (right). Additionally, as various experiments and simulations have shown [37, 38], there may be subtle differences between the motion of blood cells and that of the fluid plasma itself. A promising alternative technique involves injecting < 500 nm diameter fluorescent tracer beads into the bloodstream to allow direct high-resolution measurement of the flow of the plasma itself, even in close proximity to the vessel wall. In future work we anticipate to refine our blood flow measurements in the heart by using this tracer strategy to study in detail the relationship between the motion of blood cells and that of the surrounding plasma, and the potential impact of this phenomenon on the measured wall shear stress.

## 4. Conclusion

We have demonstrated a method for mapping the 3C-3D blood flow using z-stacks of SPIM images acquired at mutually-rotated sample orientations with multi-view data fusion. A key technical challenge in such methods is accurate co-registration of raw image datasets (of stochastically-seeded blood cells) that do not contain natural registration features. To overcome this we introduced two novel innovations: a bootstrapping strategy to determine uncertainties in PIV measurements, and a registration refinement step where consistency between the downstream processed velocity vectors themselves is used as the metric for registration.

In the blood flow vector fields we recovered using our technique, we observed a significant velocity component in the direction perpendicular to the imaging plane, which would have been overlooked using two-component imaging techniques. In particular, the blood flow near the AVC and the sinoatrial node shows a high out-of-plane velocity. This component contributes to the fluid-structure interactions, namely the wall shear stress, between the blood flow and the cardiac endothelium.

Our computational approach, achieving full 3C-3D measurements from a single imaging channel, bypasses the complexity and practical limitations of multi-camera stereoscopic imaging. By exploiting the periodicity of the cardiac blood flow to enable multiple views to be acquired sequentially, we bring 3C-3D blood flow mapping within the reach of any microscope imaging system with depth-sectioning capabilities. This ability to make accurate measurements of the 3C-3D blood flow within the heart has the potential to bring important new insights into fluid-structure interactions within the cardiovascular system during cardiac development and remodelling.

## Supporting information

Visualisation 1

Visualisation 2

## Acknowledgement

We thank Carl Tucker, Paul Strachan and their team at Bioresearch & Veterinary Services, University of Edinburgh for zebrafish husbandry and advice. KR is supported by a PhD studentship from Medical Research Scotland (PHD-50442-2021). Animals were maintained according to the Animals (Scientific Procedures) Act 1986, United Kingdom.

## Data availability

Data underlying the results presented in this paper are available in the University of Glasgow data repository [27]; analysis code is available on GitHub [28].

